# A ribosome assembly stress response regulates transcription to maintain proteome homeostasis

**DOI:** 10.1101/512665

**Authors:** Benjamin Albert, Isabelle C. Kos-Braun, Anthony Henras, Christophe Dez, Maria Paula Rueda, Xu Zhang, Olivier Gadal, Martin Kos, David Shore

## Abstract

Ribosome biogenesis is a complex and energy-demanding process requiring tight coordination of ribosomal RNA (rRNA) and ribosomal protein (RP) production. Alteration of any step in this process may impact growth by leading to proteotoxic stress. Although the transcription factor Hsf1 has emerged as a central regulator of proteostasis, how its activity is coordinated with ribosome biogenesis is unknown. Here we show that arrest of ribosome biogenesis in the budding yeast *S. cerevisiae* triggers rapid activation of a highly specific stress pathway that coordinately up-regulates Hsf1 target genes and down-regulates RP genes. Activation of Hsf1 target genes requires neo-synthesis of RPs, which accumulate in an insoluble fraction, leading to sequestration of the RP transcriptional activator Ifh1. Our data suggest that levels of newly-synthetized RPs, imported into the nucleus but not yet assembled into ribosomes, work to continuously balance Hsf1 and Ifh1 activity, thus guarding against proteotoxic stress during ribosome assembly.

## Introduction

Ribosome assembly is the most energy demanding process linked to cell growth and requires coordinated production of processed ribosomal RNAs (rRNAs), ribosomal proteins (RPs) and ribosome biogenesis (RiBi) factors. This massive biosynthetic program permits rapidly growing yeast cells to produce about 2,000 ribosomes per minutes (Warner, 1999). Rapid ribosome biogenesis is critical for sustaining high rates of growth (mass accumulation) and proliferation. Nevertheless, ribosome assembly poses a constant threat to cellular protein homeostasis since it requires the coordinated and large-scale assembly of four rRNAs with 79 different RPs, the latter of which are known to be highly prone to aggregation (David et al., 2010; Pillet et al., 2017; Rand and Grant, 2006; Weids et al., 2016). Although p53 activation upon accumulation of unassembled RPs in metazoans was first described over 20 years ago, conserved p53-independent pathways that respond to perturbations in ribosome assembly are now beginning to emerge (James et al., 2014). Given the absence of p53 in yeasts, *S. cerevisiae* promises to be a good model system in which to uncover ancestral processes that regulate growth and protein homeostasis in eukaryotes.

Heat shock factor 1 (Hsf1) is a central actor in Protein Quality Control (PQC) and protein homeostasis (proteostasis) in eukaryotes, in both stressed and unstressed cell, and in pathological situations (Li et al., 2017). Notably, Hsf1 is a direct modulator of tumorigenesis and becomes essential, as it is in budding yeast (Solis et al., 2016), to support growth of malignant cells (Santagata et al., 2013). Hsf1 prevents protein aggregation and proteome imbalance by driving the expression of a small regulon including genes encoding essential chaperones (Hsp70/Hsp90), nuclear/cytoplasmic aggregases, and proteasome components (Mahat et al., 2016; Pincus et al., 2018; Solis et al., 2016). Interestingly, studies in budding yeast reveal that the Ribosome Quality Control complex (RQC), conserved from yeast to human (Brandman et al., 2012), increases Hsf1 activity under conditions of translation stress. However, many essential aspects of Hsf1 regulation remain to be elucidated, in particular whether its transcriptional activity is linked to ribosome biogenesis itself. Recently, a conserved PQC mechanism referred to as Excess Ribosomal Protein Quality Control (ERISQ) was described that specifically recognizes unassembled RPs in the nucleus and targets them for proteasome degradation (Sung et al., 2016a; Sung et al., 2016b), thus illuminating observations made 40 years ago showing that excess RPs are rapidly degraded (Gorenstein and Warner, 1977; Warner, 1977). The potential role of Hsf1 in ERISQ has not yet been explored.

Given the tremendous investment of cellular resources involved in ribosome production (Warner, 1999) and the fact that a decrease of ribosome abundance protects cells against proteotoxic stress (Guerra-Moreno et al., 2015; Mills and Green, 2017) it might be expected that cells have evolved mechanisms to rapidly decrease RP gene transcription in the face of defects in ribosome assembly in order to both save energy and reestablish cell homeostasis. In *S. cerevisiae*, RP gene transcription is known to be tightly regulated according to growth conditions through the stress-sensitive transcription factor (TF) Ifh1. For example, Ifh1 is rapidly released from RP promoters only minutes following inhibition of the conserved eukaryotic growth regulator Target Of Rapamycin Complex 1 (TORC1) kinase (Schawalder et al., 2004). Although it has been shown that Ifh1 promoter binding is coordinated with RNA polymerase I (RNAPI) activity upon prolonged TORC1 inhibition to help balance RP and rRNA production (Albert et al., 2016; Rudra et al., 2007), how Ifh1 is removed from RP gene promoters to immediately down-regulate their expression following stress remains a mystery. Furthermore, possible links between RP gene expression, ribosomal assembly and the protein homeostasis transcription program driven by Hsf1 remain important open questions.

In this study we uncover a novel regulatory pathway, hereafter referred to as the Ribosomal Assembly Stress Response (RAStR), that allows rapid and simultaneous up-regulation of protein homeostasis genes and down-regulation of RP genes following disruption of various steps in ribosome biogenesis (rRNA production, processing or RP assembly). We show that RAStR is highly specific to the RP and Hsf1 regulons, with little or no effect on a much larger group of genes implicated in the Environmental Stress Response (ESR). Importantly, RAStR requires neo-synthesis of RPs following stress and is linked to the accumulation of RP aggregates, which we propose lead to Hsf1 activation, through chaperone competition, and to the sequestration of Ifh1 in an insoluble nucleolar fraction. Notably, we show that cycloheximide treatment leads to a transcriptional response opposite to that of RAStR, supporting a model in which unstressed cells constantly monitor nuclear levels of unassembled RPs and use this information to balance expression of Hsf1 target genes with those encoding RPs. Finally, we demonstrate that RAStR is the initial transcriptional response to inactivation of TORC1 kinase, supporting a key role for this regulatory pathway in the activation of a protein homeostasis transcriptional program that allows cells to cope with the proteotoxic consequences of disruptions to ribosome biogenesis.

## Results

### Topoisomerase depletion triggers a rapid repression of RP genes and activation of proteostasis genes

In an effort to understand the role of the two major eukaryotic DNA topoisomerases in protein-coding gene transcription, we generated yeast strains in which Top1, Top2, or both of these enzymes are rapidly degraded by the auxin-induced degron (AID) method ((Nishimura et al., 2009) and confirmed by Western blotting that significant depletion of either protein was obtained between 10 and 20 minutes following auxin addition to the medium, and that Top2 depletion, as expected, prevents cell growth (**Figure S1A, B; Table S1**). We then performed ChIP-seq analysis of RNA polymerase II (RNAPII) in the Top1-AID, Top2-AID and Top1/2-AID strains at 20 and 60 minutes following auxin addition (**Figure 1A-C**). As expected (Brill et al., 1987; Brill and Sternglanz, 1988), the absence of Top2 had little or no effect on RNAPII distribution (**Figure 1B**). However, Top1 depletion triggered a rapid response at two specific groups of genes: up-regulation of Hsf1 target genes and down-regulation of ribosomal protein (RP) genes (**Fig 1A, D, S1C-D**). Remarkably, this response was transient, as both groups of genes returned to normal levels (i.e. before auxin addition) by 60 minutes. This re-equilibration was dependent upon Top2 since it failed to occur in the Top1/2 strain, where prolonged auxin treatment led to significant dysregulation of many other RNAPII-transcribed genes (**Figure 1C**).

**Figure 1:**
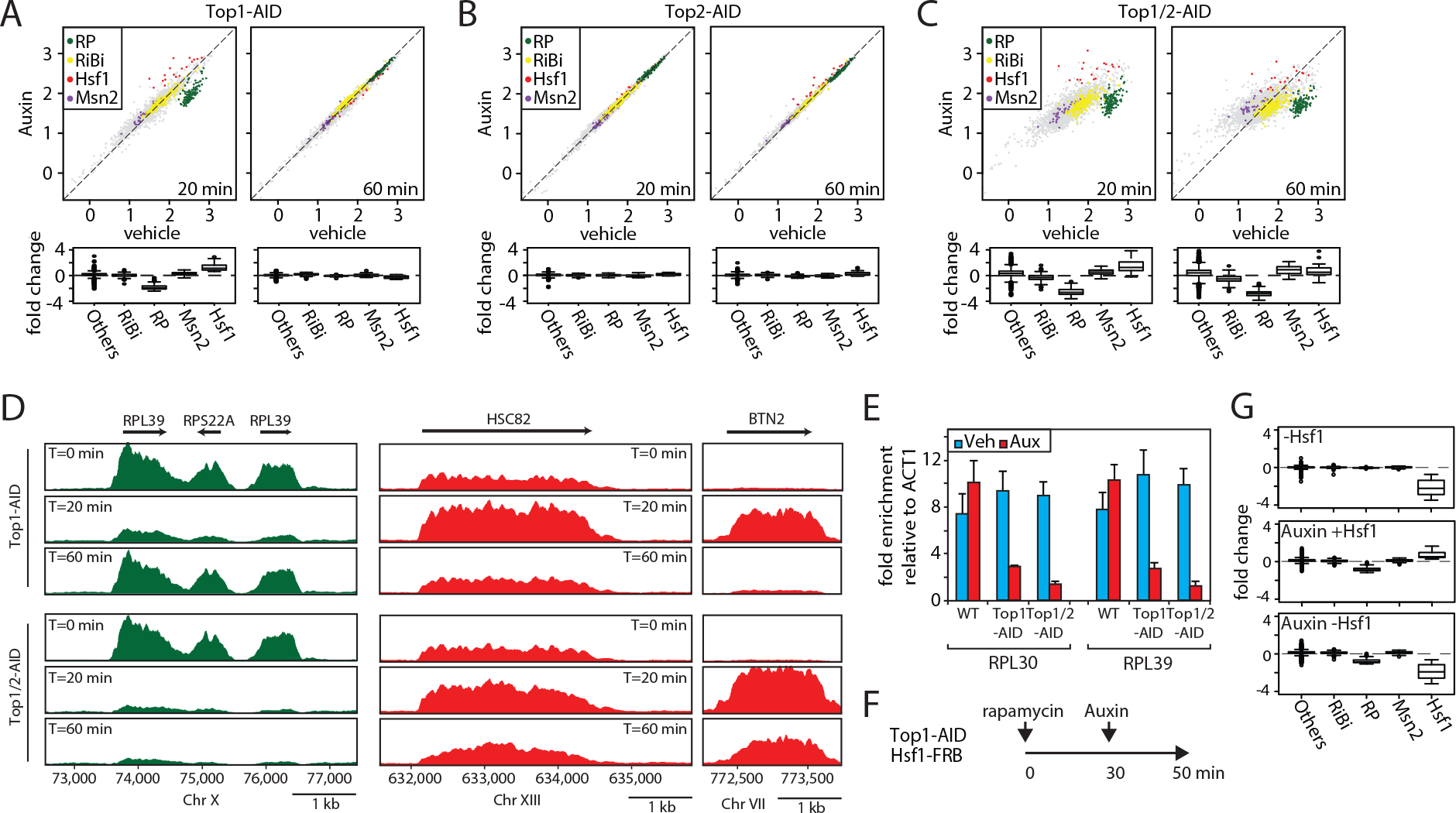
Rapid degradation of Topoisomerase 1 (Top1) induces a transient induction of Heat Shock Factor 1 (Hsf1) target genes and down-regulation of ribosomal protein (RP) genes. (**A, B, C**) Scatter plots (top panels) and box plots (bottom panels) comparing RNA polymerase II (RNAPII) binding as measured by ChIP-seq in Top1-AID (A), Top2-AID (B) and Top1/2-AID (C) strains at the indicated times following either auxin or vehicle addition to the media. Indicated gene categories (RP; ribosome biogenesis, RiBi; Msn2 target genes, and Hsf1 target genes) are color-coded on the scatter plots and displayed separately on the box plots, together with all remaining genes as a fifth class (other). (**D**) Genome browser tracks showing RNAPII ChIP-seq read counts at the indicated positions of chromosomes X, XIII and VII at 0, 20, or 60 minutes following auxin addition to Top1-AID (top panels) and Top1/2-AID (bottom panel) strains. Gene names and open reading frame (ORF) positions are shown above. (**E**) Ifh1 occupancy, measured by qPCR ChIP at the RPL30 and RPL30 promoters 20 minutes following auxin addition to cultures of Top1-AID and Top1/2-AID strains. (**F**) Schematic representation of protocol for Hsf1-FRB nuclear depletion (anchor-away) followed by Top1-AID depletion (auxin-induced degron). (**G**) Box plots showing RNAPII ChIP-seq signal following Hsf1-FRB nuclear depletion by anchor-away (rapamycin, top panel), Top1-AID degradation (auxin, middle panel) or both Hsf1-FRB and Top1-AID depletion (rapamycin + auxin, bottom panel) for the five indicated categories of genes.

Since up-regulation of proteostasis-related genes and down-regulation of RP genes are characteristic of many different stress responses, we decided to quantify the effect of Top1 depletion on transcription all gene groups that have been classified as part of the general “Environmental Stress Response” (ESR; (Gasch et al., 2000)), which include an additional group of stress-induced genes regulated by the Msn2/4 TFs and a large suite of genes involved in ribosome biogenesis (RiBi genes). This analysis shows clearly that Top1 depletion, as well as depletion of both Top1 and Top2 (at the early 20-minute time point), triggers a highly specific stress response linked to RP genes and Hsf1 target genes (**Figure 1A-C**). Such a targeted response is unlikely to result from a global topological effect on RNAPII recruitment but would instead appear to be the consequence of the activation of a specific signaling pathway that is more restricted in nature than the ESR.

To explore the target(s) of this hypothetical signaling pathway at RP genes we monitored by qPCR ChIP the promoter association of three TFs (Rap1, Fhl1 and Ifh1) that operate at the majority (>90%) of these 138 genes (Knight et al., 2014). Interestingly, we found that the activator Ifh1 is rapidly released from RP gene promoters after topoisomerases depletion (**Figure 1E**) whereas Rap1 and Fhl1, which bind directly to RP promoter DNA, are not affected (**Figure S1E-F**). To confirm that Hsf1 is indeed required for up-regulation of genes following Top1 depletion we used the anchor-away technique (Haruki et al., 2008) to rapidly remove Hsf1 from the nucleus before initiating Top1-AID degradation (**Figure 1F**). This revealed that Hsf1 nuclear depletion completely abolishes activation of stress genes following Top1 depletion without affecting down-regulation of RP genes (**Figure 1G**). Therefore, activation of stress-induced genes following Top1 depletion is completely Hsf1-dependent, whereas repression of RP genes is independent of Hsf1 or induction of its target genes. We would also note that the stress pathway induced by Top1 depletion is unusually restricted in comparison to many other stress responses that are often grouped together as the Environmental Stress Response (ESR; (Gasch et al., 2000)), since Msn2/4 target genes are not induced and Ribi genes are not down-regulated (**Figure 1A, G**).

### Top1 depletion arrests ribosome biogenesis and activates a ribosomal assembly stress response

Although it may seem surprising that depletion of topoisomerases can induce a Hsf1-dependent stress response, formation of distinct nuclear foci by the Btn2 aggregase and perinuclear accumulation of the proteasome subunit Pre6 following Top1/2 degradation (**Figure S2A, B**) both point to the induction of proteotoxic stress in the nucleus (Miller et al., 2015a). Top1 was initially identified through a mutant (*MAK1*) defective in large ribosomal subunit production (Thrash et al., 1985) and later shown to be required for proper rRNA synthesis (Brill et al., 1987; El Hage et al., 2010; French et al., 2011). Consistent with these findings, we observed a strong reduction of pre-rRNA synthesis as shown by decreased labelling of the RNAPI-transcribed 35S pre-rRNA, and two co-transcriptionally cleaved products, 27S and 20S pre-rRNAs, only 10 minutes after Top1 (or Top1/2) depletion (**Figure S2C**). This decreased synthesis is concomitant with an elongation defect, as shown by the accumulation of truncated pre-rRNA, initially described by Tollervey lab (El Hage et al., 2010), in the absence of Top1 (**Figure 2A**). This rapid defect in rRNA production by inhibition of RNAPI elongation leads to unbalanced production of 40S and 60S ribosomal subunits, with a marked deficiency of the large (60S) subunit relative to the small (40S) subunit (**Figure 2B**). This would be expected to create a disequilibrium between RP and rRNA production, and more specifically an excess of unassembled RPs. Consistent with this, we detect accumulation of a large subunit protein, Rpl3, in trailing fractions of polysome gradients (**Figure 2C**). These observations strongly suggest that RPs fail to be incorporated normally into ribosomes immediately following Top1 degradation. RPs are known to be prone to aggregation (David et al., 2010; Pillet et al., 2017; Rand and Grant, 2006; Weids et al., 2016) and recent reports show that newly-synthetized, unassembled RPs accumulate in aggregates in response to ribosome assembly stress (Sung et al., 2016a; Sung et al., 2016b). Significantly, we also observed accumulation of RPs in an insoluble fraction following topoisomerase degradation (**Figure 2D**).

**Figure 2:**
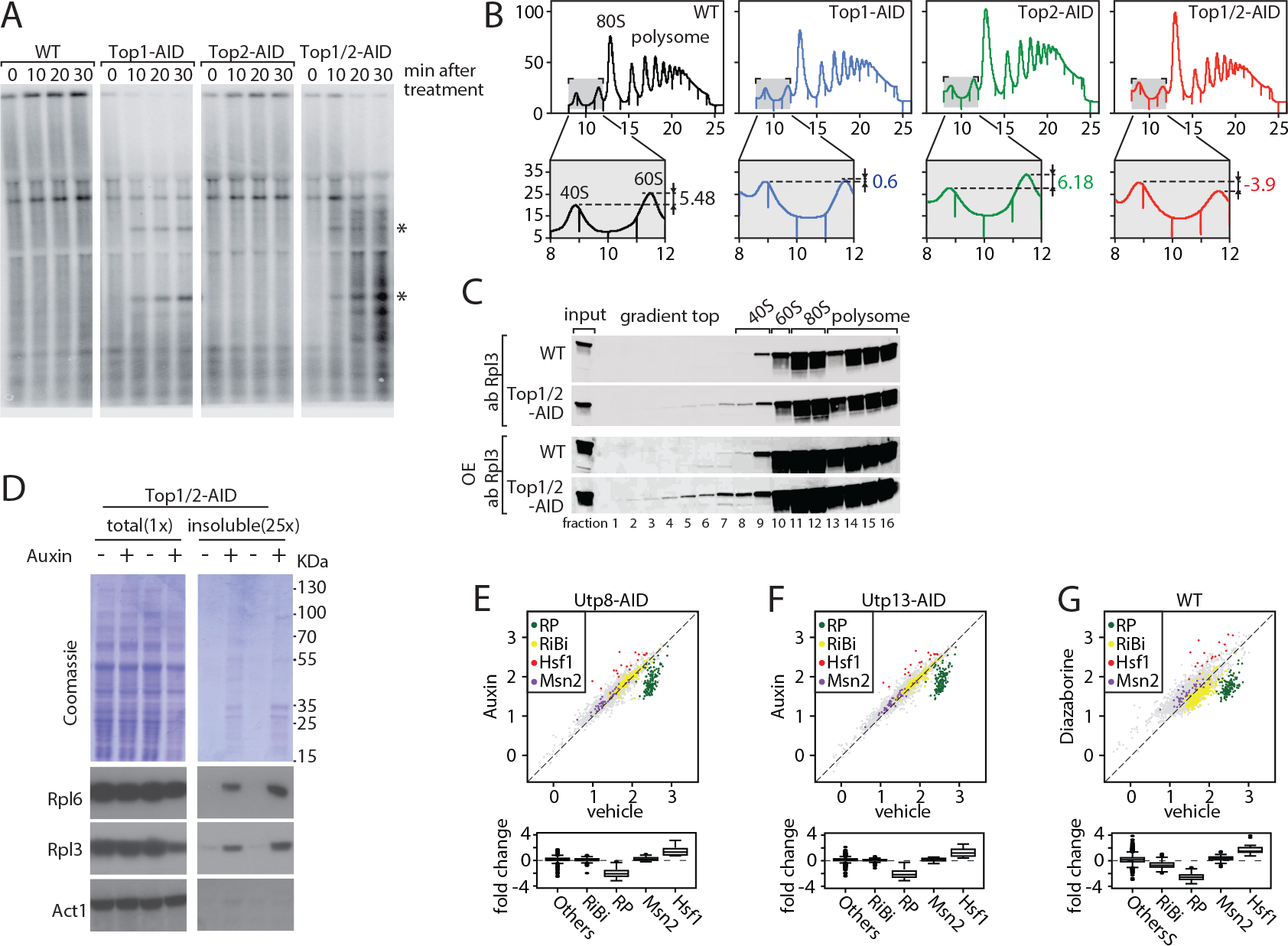
Evidence linking the effect of Top1 depletion on RNAPII to an underlying defect in ribosome biogenesis. (**A**) Northern blot of RNAs prepared from cultures of wild-type, Top1-AID, Top2-AID and Top1/2-AID cells at the indicated times following addition of auxin to the media. Total RNAs were extracted and samples were separated on agarose gels, transferred to a nylon membrane and hybridized with specific oligonucleotide probes. Truncated pre-rRNA fragments that accumulate in the absence of Top1 (Top1-AID) or both Top1 and Top2 (Top1/2-AID) are indicated (*). (**B**) Polysome sedimentation profiles (OD260) of WT, Top1-AID, Top2-AID, and Top1/2-AID strains 20 minutes following auxin treatment (top panels). The top of each gradient, corresponding to 40S and 60S subunit peaks is expanded below, where peak height differences (60S-40S) are indicated. (**C**) Total cell extracts prepared from the indicated fractions of sedimentation profiles of WT and Top1/2-AID strains in (B), TCA precipitated and analyzed by western blot following SDS-PAGE, using an antibody against Rpl3. (**D**) Total (left panels) and detergent-insoluble pellet (right panels) fractions isolated from lysates of Top1/2-AID cells treated (+) or not (−) with auxin were analyzed by SDS-PAGE and Coomassie blue staining (top panels) or immunoblotting with the indicated antibodies (bottom panels). The pellet fraction is overloaded 25-fold compared to the total fractions. (**E-F-G**) Scatter plots (top panels) comparing RNAPII ChIP-Seq read counts for individual genes in Utp8-AID (E) or Utp13-AID (F) cells after 20 minutes of auxin or vehicle treatment, or WT cells after 20 minutes of treatment with diazaborine or vehicle (G) (y-axis: auxin or diazaborine) for 20 minutes versus non-depleted cells (x-axis, Vehicle). Each dot represents a gene (5044 genes in total) and genes are color-coded according to functional groups as in Figure 1C. Bottom panels display the corresponding box plots for the four indicated gene categories plus all other genes (others).

The observations described above led us to hypothesize that the transcriptional response to Top1 degradation is a consequence of defective ribosome assembly, perhaps driven by proteotoxic stress caused by the accumulation of unassembled RPs. To begin to test this idea, we measured the transcriptional response to three different perturbations to ribosome biogenesis: depletion of two essential rRNA processing factors (Utp8 and Utp13) and treatment of cells with diazaborine, a drug that rapidly blocks 60S subunit maturation. Remarkably, all of these treatments triggered a similar transcriptional response to that which occurs following Top1 depletion, namely a specific down-regulation of RP genes and up-regulation of Hsf1 target genes (**Figure 2E, G; Table S2**), which we refer to as the “Ribosome Assembly Stress Response” (RAStR).

Hsf1 activity is stimulated by many different types of cellular stress, including stalled ribosomes. A pioneering study reported that a set of proteins termed the Ribosome Quality Control Complex (RQC) binds to 60S ribosomal subunits containing stalled polypeptides and leads to their degradation. In the process, the RQC triggers a specific stress signal that leads to Hsf1 target gene activation (Brandman et al., 2012). Thus, cells lacking a component of the RQC, the Tae2 protein, fail to activate Hsf1 following translational stress. To ask if RAStR might be related to the RQC, we induced Top1 degradation in *tae2-*Δ cells. We found that Hsf1 activation was identical to that observed in the *TAE2* parental strain (**Figure S2D**) and conclude that RAStR and the RQC activate Hsf1 through different mechanisms. These results highlight that cells have developed distinct mechanisms to adapt the Hsf1 transcriptional program to defects in both ribosome activity and ribosome assembly.

### Ifh1 is sequestered in an insoluble nuclear fraction during RAStR

Although many studies would support the notion that Hsf1 activation during RAStR occurs through sequestration of its inhibitory partner Hsp70 by RP aggregates (Krakowiak et al., 2018; Shi et al., 1998; Zheng et al., 2016), it is less clear how ribosome assembly stress could trigger release of Ifh1 from RP gene promoters. We reported previously that the association of Ifh1 with RP gene promoters in growing cells is rapidly disrupted (within 5 min) following inhibition of the growth-promoting TORC1 kinase by addition of rapamycin to the medium (Schawalder et al., 2004). More recently (Albert et al., 2016), we found that stable release of Ifh1 from RP gene promoters (measured 20 min after rapamycin addition) requires its C-terminal domain together with a complex of proteins containing casein kinase 2 (CK2) and two RiBi factors, Utp22 and Rrp7, with which Ifh1 interacts to form the CURI complex (Rudra et al., 2007). Thus, in *ifh1-*ΔC cells Ifh1 is rapidly released but later returns to RP gene promoters following TORC1 inhibition. This led us to propose two distinct mechanisms controlling the promoter release of Ifh1 following stress: one operating at a short timescale (< 5 min) and the other on a long timescale (^~^20 min). Interestingly, ifh1-ΔC release is stable following Top1 depletion, suggesting that an unknown mechanism regulates Ifh1 during RAStR (**Figure 3A**).

**Figure 3:**
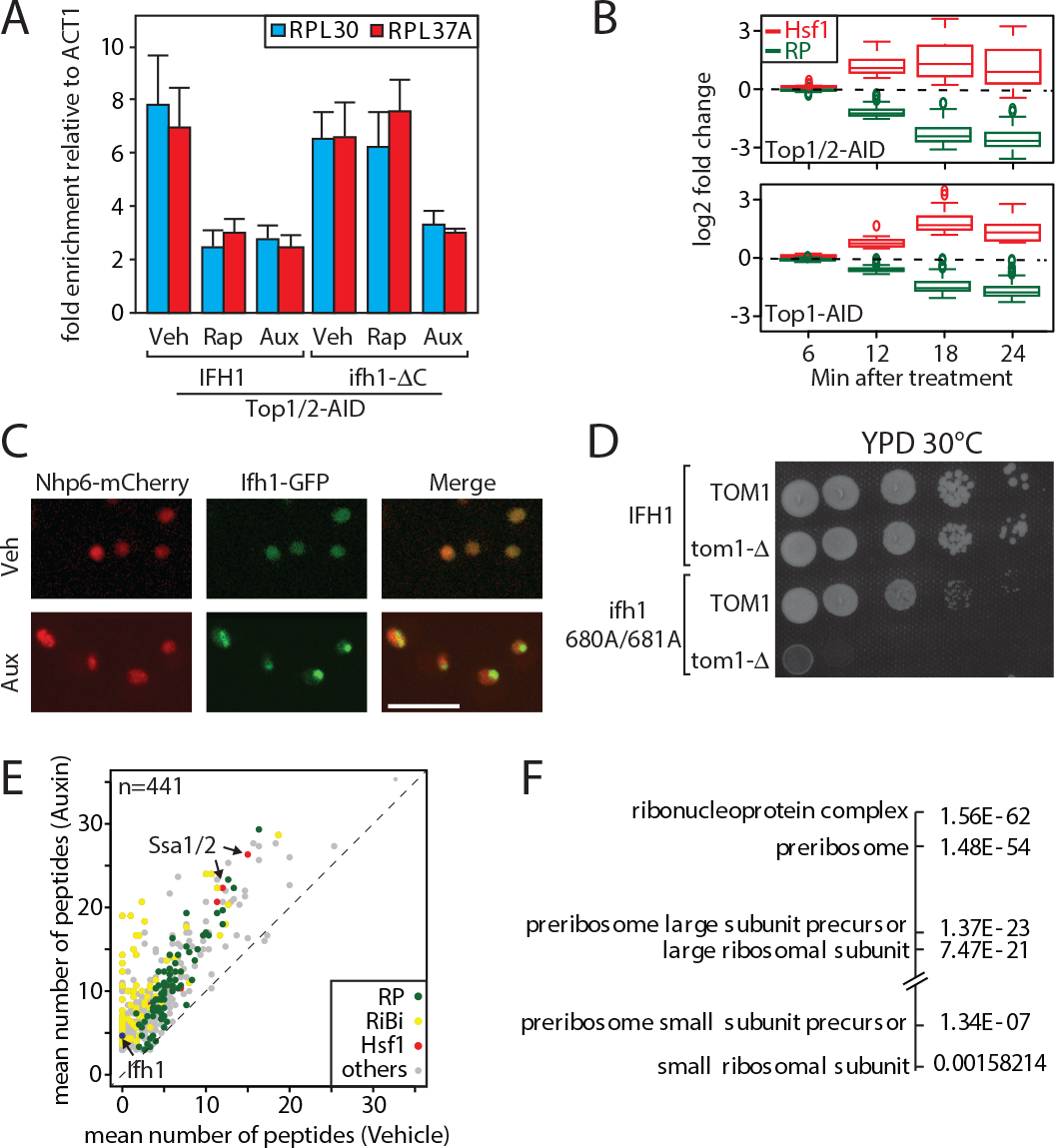
Evidence that Ifh1 is rapidly removed from RP gene promoters and sequestered in an insoluble nucleolar fraction pursuant to unassembled RP accumulation following RAStR initiation. (**A**) Ifh1 occupancy at the RPL30 and RPL39 promoters 20 minutes following vehicle, rapamycin or auxin treatment of Top1/2-AID strains expressing either WT Ifh1 (Ifh1; left panel) or a C-terminal truncated allele (*ifh1*-1ΔC; right panel). (**B**) Box plots showing the kinetics of RNAPII ChIP-seq changes at Hsf1 target and RP genes at the indicated time points (minutes) following auxin treatment in Top1/2-AID (top panel) or Top1-AID (bottom panel) strains. (**C**) A Top1/2-AID strain expressing Ifh1-eGFP and mCherry-Nhp6 was grown exponentially and cell samples were used for fluorescence microscopy analysis after 20 minutes of auxin (Aux) or vehicle (Veh) treatment. (**D**) 10-fold serial dilutions of IFH1 or ifh1-AA cells transformed in either *TOM1* of *tom1*-Δ backgrounds (as indicated) were grown in YPAD medium for 44 hours at 30°C before being photographed. (**E**) Scatter plot comparing average number of peptides purified in an insoluble fraction from Top1/2-AID cells treated for 20 minutes with either auxin (y-axis, Auxin) or vehicle (x-axis, Vehicle). Each dot represents a protein, color-coded according to functional group as above (green: RP, red: Hsf1 target gene product, yellow: RiBi protein, grey: others), with some specific proteins indicated by arrows (n=3). (**F**) Gene Ontology and p-values of protein groups that are the most enriched in the insoluble fraction following Top1/2-AID depletion (Δ>3 peptides in insoluble fraction after topoisomerase depletion compared to vehicle in all experiments, n=3).

The fact that RP gene repression and Hsf1 target gene activation occur with identical kinetics following RAStR activation (**Figure 3B**), and that Ifh1 concentrates in nuclear foci rapidly after topoisomerase depletion (**Figure 3C**), suggests that Ifh1 could be sensitive to the accumulation of unassembled RPs in the nucleus, as is presumably the case for Hsf1. To assess this possibility, we first analyzed published mass spectrometry data of insoluble fractions from cells treated with a proteasome inhibitor or lacking Tom1, an E3 ligase required for degradation of unassembled RPs (Sung et al., 2016a; Sung et al., 2016b). In both cases RPs accumulate in an insoluble fraction, together with chaperones, including Hsp70, the inhibitory partner of Hsf1. Interestingly, Ifh1 is also strongly and specifically enriched in the presence of RP aggregates (**Figure S3A**) indicating that Ifh1 could be trapped in an insoluble cellular fraction in the absence of Tom1 and thus unable to efficiently associate with RP gene promoters. To test this possibility, we combined deletion of *TOM1* with a mutant allele of *IFH1* (*ifh1-AA*) that weakens its interaction with RP gene promoters. Remarkably, *tom1-*Δ is synthetically lethal with *ifh1-AA* (**Figure 3D**) supporting the notion that RP aggregation could directly impact on Ifh1 promoter binding.

To assess directly whether Ifh1 is sequestered in aggregates during RAStR, we analyzed by mass spectrometry the insoluble fraction following topoisomerase depletion. As previously reported for *tom1-*Δ cells (Sung et al., 2016a), the insoluble fraction is enriched in chaperones and RPs (**Figure 3E, F**). However, we also note a strong increase of RiBi factors, primarily those implicated in biogenesis of the large ribosomal subunit (**Figure 3F**). Importantly, Ifh1 was never detected in an insoluble fraction in the absence of stress but was invariably detected in these fractions following topoisomerase depletion (**Table S3**). We argue that this rapid sequestration of Ifh1 may be sufficient to explain the observed down-regulation of RP genes during RAStR.

### Neo-synthetized RPs are required for RAStR activation

Given their fast turnover rate, nuclear accumulation in the absence of ribosome assembly and propensity to aggregate, unassembled RPs could be ideally positioned to rapidly signal ribosome biogenesis defects (Milkereit et al., 2001; Sung et al., 2016a). To evaluate the importance of newly synthetized RPs in RAStR, we blocked their production by cytoplasmic anchoring of Ifh1 before topoisomerase depletion (**Figure 4A**). It is important to note that Ifh1 binding is highly specific to RP genes (Knight et al., 2014) and that the transcriptional effect of its nuclear depletion is restricted to RP genes and a very small number of additional targets. We note that although Ifh1 depletion by anchor-away is probably not complete (the strain still grows, albeit slowly, even though Ifh1 is essential for growth; **Figure S4A**), it nonetheless does lead to a significant and highly specific decrease in RP gene transcription as measured by RNAPII ChIP-seq (**Figure 4B; Table S4**). Interestingly, Ifh1 depletion also leads to aberrant rRNA processing (**Figure 4C**) as would expected in conditions where RP levels become limiting (Reiter et al., 2011). Remarkably, we noted that Ifh1 anchoring alone, in the absence of topoisomerase depletion, also caused a significant down-regulation of Hsf1 target genes (**Figure 4D; Table S4**) even though Ifh1 is absent from the promoters of these genes (**Figure S4B**), suggesting that the Hsf1 transcriptional program is continuously influenced by RP production. Consistent with this idea, we found that Top1 up-regulation of Hsf1 genes was either abolished or strongly reduced (**Figure 4E, F, G; Table S4**) when Top1 or Top1 and Top2 were degraded following nuclear depletion of Ifh1, indicating that RP production is required for Hsf1 target gene activation during RAStR.

**Figure 4:**
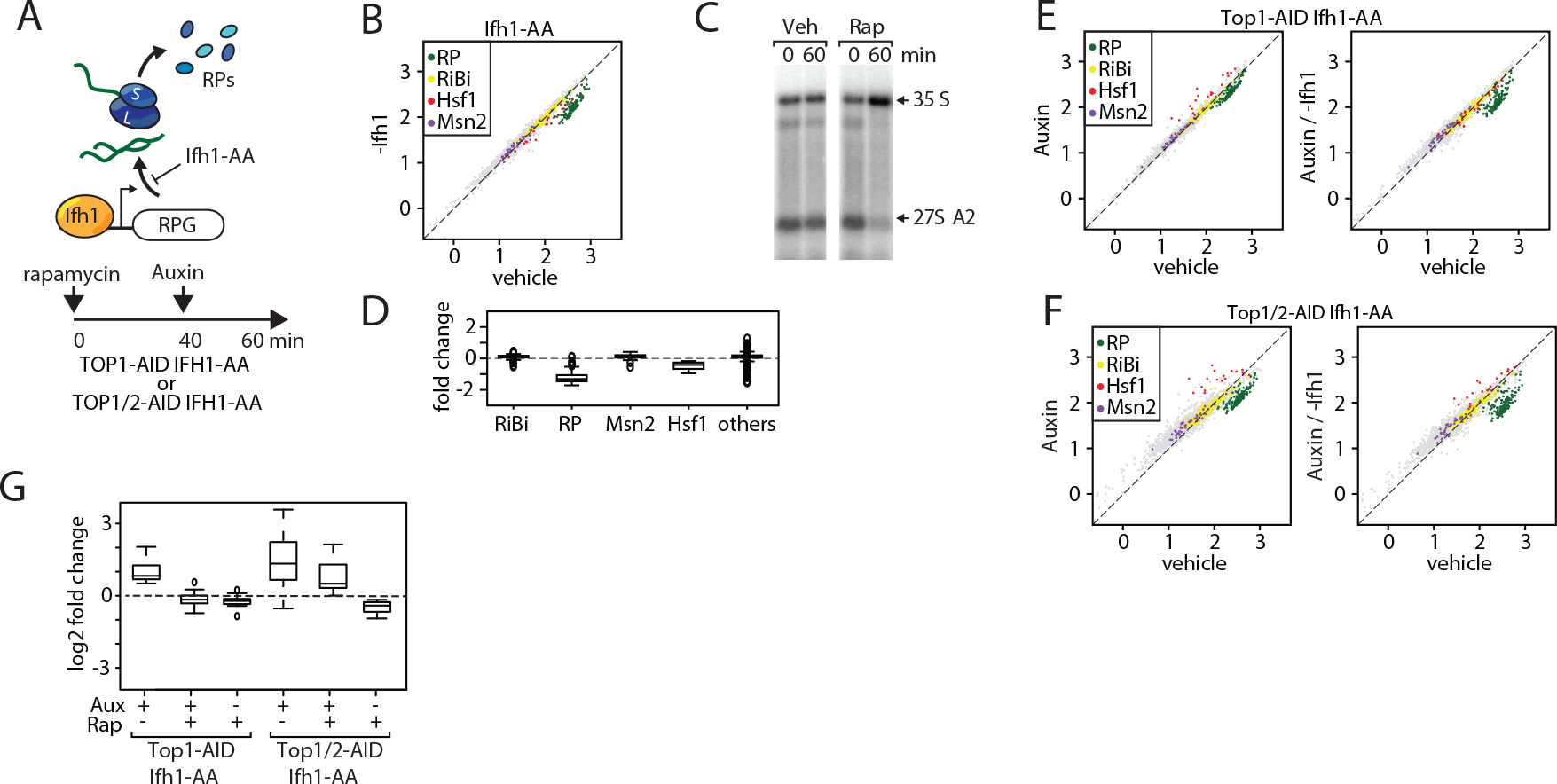
Down-regulation of RP gene expression by Ifh1 nuclear depletion prior to RAStR initiation strongly dampens Hsf1 target gene activation. (**A**) Schematic of protocol for Ifh1-FRB nuclear depletion (0-60 minutes of rapamycin treatment) followed by Top1-AID or Top1/2-AID degradation (auxin treatment, 40-60 min). (**B**) Scatter plot comparing RNAPII ChIP-Seq in Ifh1-FRB cells either rapamycin-treated for 60 min (y-axis, Rap, Ifh1-FRB nuclear depletion) or un-treated (x-axis, Veh, no Ifh1-FRB depletion). Categorization and color-coding of genes as above. (**C**) Northern blots of pre-rRNA after 0 or 60 minutes of vehicle (Veh) or rapamycin (Rap) treatment of Ifh1-FRB strain. (**D**) Box plots of the data shown in (B) for the indicated five gene categories, showing fold-change upon Ifh1-FRB nuclear depletion compared to mock treated cells. (**E**) Scatter plots comparing RNAPII ChIP-Seq in Top1-AID Ifh1-FRB cells either auxin-treated (y-axis, Aux, left panel) or auxin-plus rapamycin-treated (y-axis, Aux / -Ifh1, right panel) treated, as described in (A), versus untreated cells (x-axis, Vehicle, both panels). (**F**) As in (E), but for Top1/2-AID Ifh1-FRB cells. (**G**) Box plots showing RNAPII ChIP-seq change after rapamycin and/or auxin treatment for Hsf1 target genes in Top1-AID Ifh1-FRB cells (left) or Top1/2-AID Ifh1-FRB cells (right).

### Cells balance RP production and Hsf1 activity even in the absence of ribosome assembly stress

As an alternative approach to test the requirement for *de novo* RP synthesis to initiate RAStR we used cycloheximide treatment, which is known to rapidly deplete the nuclear pools of RPs (**Figure 5A**;(Gorenstein and Warner, 1977; Lam et al., 2007; Reiter et al., 2011; Warner, 1977)). As reported by others, cycloheximide treatment alone triggers a rapid arrest of rRNA processing (**Figure 5B**). We also noted, quite strikingly, that cycloheximide treatment induces a transcriptional response exactly opposite to that induced by RAStR, namely Hsf1 target gene down-regulation and RP gene up-regulation (**Figure 5C**; **Table S5**). This finding suggests that even in unstressed cells RP production may contribute to a basal level of Hsf1 activation while at the same time limiting Ifh1 activity at RP gene promoters.

**Figure 5:**
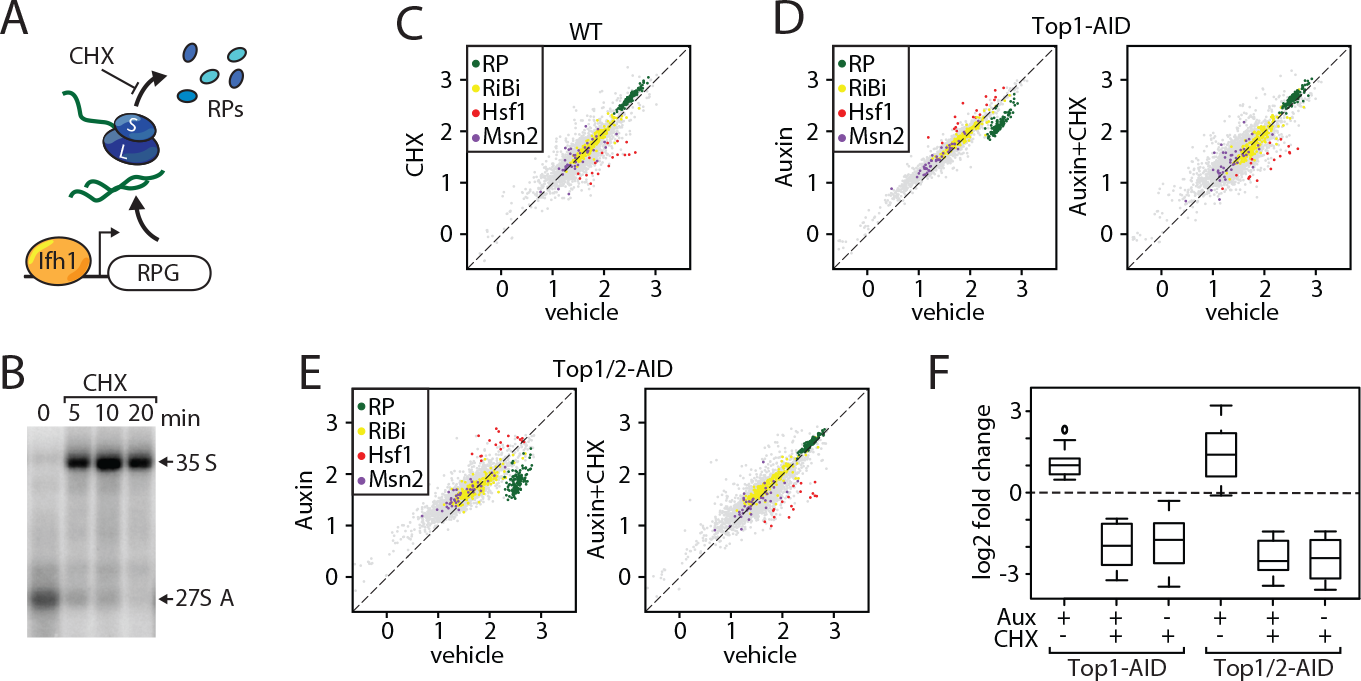
Cycloheximide treatment induces a rapid RNAPII transcriptional response opposite and epistatic to that of RAStR. (**A**) Schematic describing the effect of cycloheximide (CHX) treatment on de novo RP production. (**B**) Northern blots of pre-rRNA after 0, 5, 10, 20 minutes of cycloheximide (CHX) treatment. (**C**) Scatter plot comparing RNAPII ChIP-Seq after 20 minutes of cycloheximide treatment (y-axis, CHX) to that of non-treated cells (x-axis, Vehicle). (**D**) Scatter plots comparing RNAPII ChIP-Seq in auxin-treated Top1-AID cells, treated (y-axis, Auxin+CHX, right panel) or not (y-axis, Auxin, left panel) with cycloheximide, to untreated cells (neither auxin nor cycloheximide; x-axis, Vehicle). (**E**) Scatter plots comparing RNAPII ChIP-Seq as in (D), but for Top1/2 cells. (**F**) Box plots showing RNAPII ChIP-seq fold-change for Hsf1 target genes after cycloheximide (CHX) and/or auxin (Aux) treatment Top1-AID or Top1/2-AID

Significantly, treatment of cells undergoing topoisomerase depletion with cycloheximide completely abolished RP gene repression (**Figure 5D, E; Table S5**), release of Ifh1 from RP gene promoters (**Figure S5**) and activation of Hsf1 target genes (**Figure 5D, E, F; Table S5**). These findings clearly demonstrate that RAStR is dependent upon *de novo* protein synthesis. Taken together with the effect of Ifh1 cytoplasmic anchoring on Hsf1 target gene activation, both in the presence and absence of topoisomerase degradation, our observations on the effect of cycloheximide highlight the interwoven nature of RP and Hsf1 target gene regulons and further support the notion that unassembled, aggregated RPs constitute the primary RAStR-induced signal capable of regulating both Ifh1 and Hsf1 activities, albeit in an opposite direction. More generally, these data indicate that newly synthetized RPs, in both stressed and unstressed cells, operate as a central hub in coordinating the expression of RP genes themselves with the Hsf1-dependent activation of chaperone and proteasome genes.

### RAStR is the first transcriptional response to environmental stress

We next turned our attention to the potential involvement of RAStR during more general stress responses that might also rapidly affect ribosome assembly. To this end, we inactivated the conserved eukaryotic growth-promoting TORC1 kinase by treatment of cells with rapamycin, which is known to mimic a major part of the environmental stress response, including high salt, redox stress, and carbon, nitrogen, phosphate or amino acid starvation (Loewith and Hall, 2011). As reported previously, rapamycin triggers a rapid arrest of rRNA processing (**Figure 6A**) and a decrease of RP and RiBi gene expression (**Figure 6B; Table S6**). Interestingly, we noted that Hsf1 target genes are transiently up- and down-regulated at 5 and 20 minutes, respectively, (**Figure 6B; Table S6**) suggesting that RAStR could be activated specifically at the early time point but not later. Consistent with this view, it has been reported that RP production ceases around 15 minutes after rapamycin treatment (Reiter et al., 2011), which we suggest would turn off the signal for RAStR, thus explaining the down-regulation of Hsf1 target genes observed at 20 minutes. Taking advantage of our observation that cycloheximide treatment abolishes activation of RAStR, we treated cells with cycloheximide 5 min before rapamycin treatment (**Figure 6C**). Remarkably, this specifically prevented RP gene repression and Hsf1 target gene activation at 5 minutes following rapamycin addition, whereas at the longer time point RP genes and Hsf1 were regulated independently of cycloheximide (repressed and activated, respectively; **Figure 6D; Table S6**). Consistent with the block in RP gene down-regulation immediately following rapamycin addition, we showed that cycloheximide treatment prevents release of Ifh1 at 5 minutes, but not at 20 minutes (**Figure 6E**).

**Figure 6:**
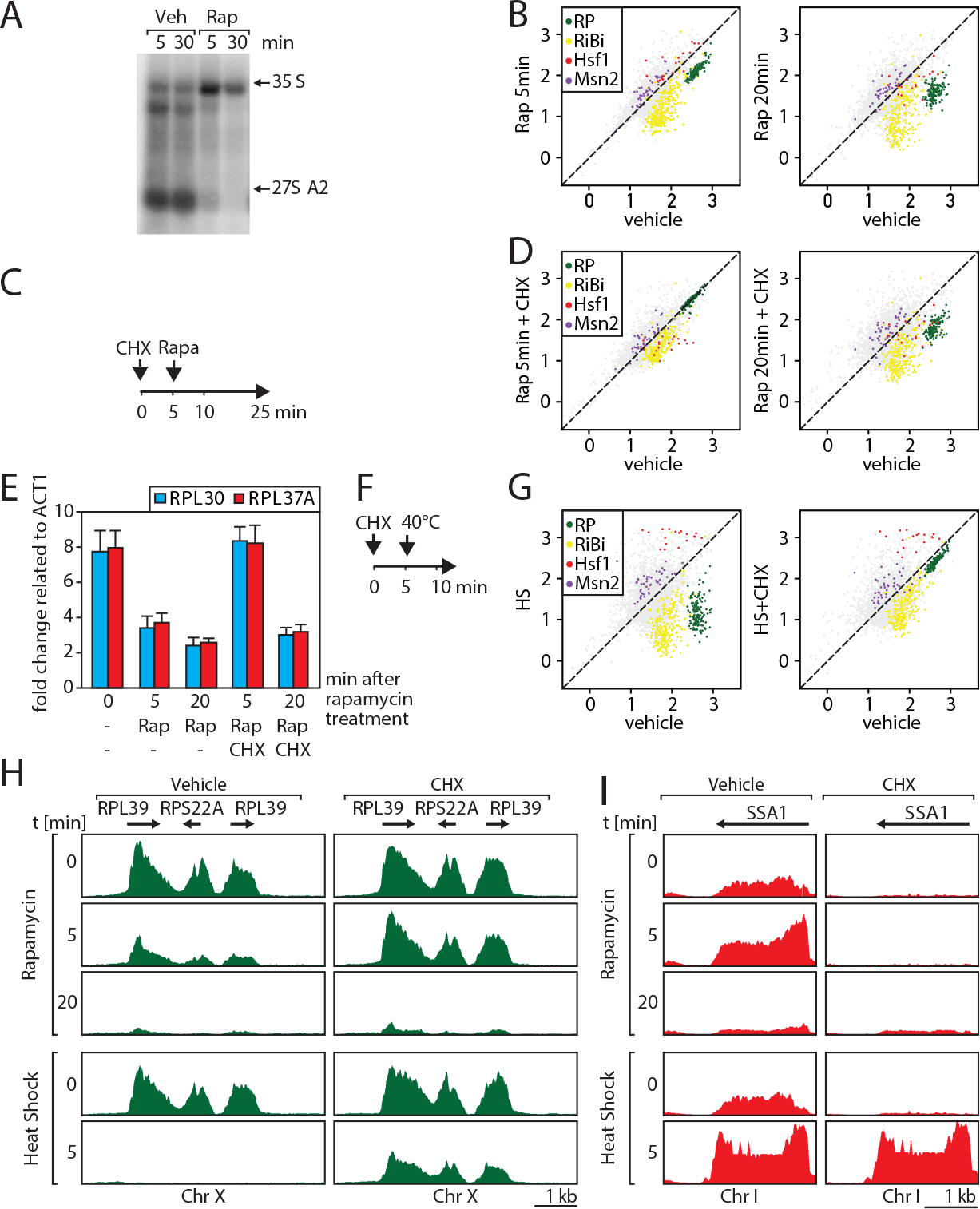
RAStR comprises the cycloheximide-sensitive component of the early RNAPII response to TORC1 inhibition. (**A**) Northern blots of pre-rRNA from WT cells after 5 or 30 minutes of vehicle (Veh) or rapamycin (Rap) treatment. (**B**) Scatter plots comparing RNAPII ChIP-Seq in cells after 5 (y-axis, Rap 5 min; left panel) or 20 minutes (y-axis, Rap 20 min, right panel) of rapamycin treatment to non-treated cells (x-axis, Vehicle, both panels). Gene groups are color-coded as before. (**C**) Schematic representation of experiment in (D): WT cells are pre-treated with cycloheximide for 5 minutes (0-25 min) before rapamycin addition (Rap, 5-25 min). Cells are collected after 5 or 20 minutes of rapamycin treatment for RNAPII ChIP-seq analysis. (**D**) Scatter plots comparing RNAPII ChIP-Seq in cells pre-treated with cycloheximide (CHX) then treated for 5 (y-axis, Rap 5min+CHX; left panel) or 20 (y-axis, Rap 20min+CHX; right panel) minutes with rapamycin versus untreated cells (x-axis, Vehicle). (**E**) Ifh1 occupancy at the RPL30 and RPL37A promoters following 5 or 20 minutes of rapamycin treatment (Rap) in cells pre-treated or not with cycloheximide (CHX) for 5 minutes. (**F**) Schematic of protocol for heat shock treatment (HS 40°C, 5-10 min) following a 5-minute CHX pre-treatment (CHX, 0-15 min) or mock treatment. Samples for ChIP-seq analysis of RNAPII association are taken at 15 minutes. (**G**) Scatter plots comparing RNAPII ChIP-Seq in cells pre-treated or not with cycloheximide (CHX) followed by 5 minutes of heat-shock (y-axis, HS) versus non-stressed cells (x-axis, untreated). (**H-I**) Genome browser tracks showing RNAPII ChIP-Seq read counts on a region of chromosome X containing three RP genes (H) or at the *SSA1* gene on chromosome I. Cells were either rapamycin- or heat shock-treated (top 3 panels and bottom 2 panels, respectively) for the indicated times, and either mock-treated (Vehicle; left panels) or cycloheximide-treated (CHX; right panels) as described in (C) and (F) above. Gene annotations are shown above the tracks. RP genes are in green (H), and the Hsf1 target gene *SSA1* is in red (I).

The effects of rapamycin treatment described above are fully consistent with our previous report demonstrating that regulation of RP gene transcription following TORC1 inactivation operates through two distinct mechanisms at short and long timescales, with the latter dependent on RNAPI activity (Albert et al., 2016). The short timescale mechanism, described here and mediated by RAStR, allows cells to rapidly arrest RP production and avoid or minimize proteotoxic stress induced by arrest of ribosome assembly. The second mechanism permits the resumption of RP production only when rRNA synthesis also resumes. These two mechanisms could be particularly useful to rapidly adapt ribosome production to new growth conditions.

To explore this idea further, we explored the effect of cycloheximide pre-treatment on the response to heat shock (**Figure 6F; Table S6**), which is known to transiently down-regulate both RP gene and rRNA transcription. In accord with a role for RPs in this process, we found that cycloheximide pre-treatment prevents strong and immediate repression of RP genes following heat shock (**Figure 6G-H; Table S6**), consistent again with the idea that RAStR is the first transcriptional response to environmental stress. Importantly, Hsf1 target genes are still activated even in presence of cycloheximide in the case of heat-shock, presumably because thermal stress will activate Hsf1 in the absence of new protein synthesis (**Figure 6G, I**). Taken together, these data demonstrate the existence of a new pathway allowing coordination of Hsf1 gene activation with RP gene expression (**Figure 7**), permitting a rapid adaptation of the Hsf1-driven protein homeostasis transcriptional program with ribosome assembly under various stress conditions.

**Figure 7:**
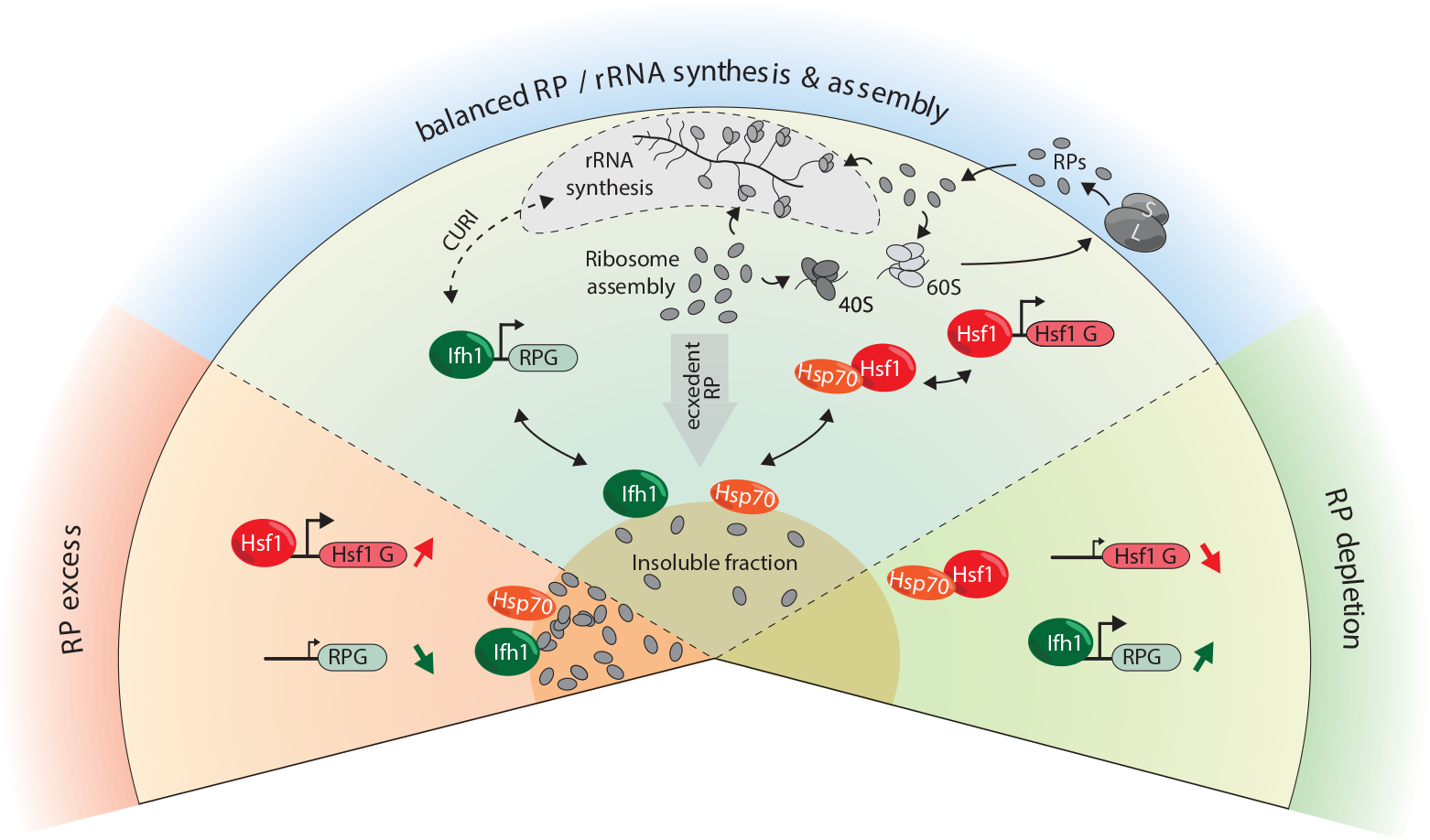
Model for RP and Hsf1 target gene regulation as a function of unassembled RP levels in both growing and stressed cells. In rapidly-growing, un-stressed cells, RNAPII initiation at RP and Hsf1 target genes is continuously adjusted according to the levels of un-assembled RPs (central sector). Under various stress conditions (including RAStR, heat shock, TORC1 inhibition, perhaps many others) levels of unassembled RPs increase dramatically, and RPs accumulate with nucleolar proteins, Ifh1 and chaperones in an insoluble nuclear or nucleolar fraction. This leads to the rapid up-regulation of Hsf1 target genes (e.g. chaperones and proteasome components), presumably through Hsp70 titration, and to the coincident down-regulation of RP genes, through Ifh1 sequestration (bottom left sector, “RP excess”). Conversely, a decrease in RP production (as provoked here by cycloheximide treatment) will lead to an opposite transcriptional response (bottom right sector, “RP depletion”). In summary, we propose that levels of unassembled nuclear RPs act to constantly adjust RP and Hsf1 target gene expression, allowing the cell to balance growth with protein homeostasis.

## Discussion

In this study we demonstrate the existence of a regulatory mechanism, which we refer to as RAStR, that allows yeast cells to specifically coordinate the activity of two TFs, Hsf1 and Ifh1, with the functional state of ribosome assembly. Taken together, our data and previous reports suggest that rapid ribosome biogenesis is a potentially proteotoxic process, in large part due to accumulation of unassembled RPs, that needs to be carefully coordinated at the transcriptional level with the production of chaperones and proteasome components (**Figure 7**). In this perspective, RAStR is essential to match ribosome production rates with the proteostasis burden that it imposes.

RPs are among the most abundant ubiquitinylated proteins that accumulate in the nucleus of proteasome-deficient *S. cerevisiae* and human cells (Lam et al., 2007; Mayor et al., 2007; Sung et al., 2016a; Sung et al., 2016b), suggesting that the synthesis of RPs and their assembly into ribosomes must be tightly coordinated with the cell’s proteostasis capacity. Consistent with this view, we show here that induction of ribosome assembly stress is correlated with the rapid accumulation of RPs in a detergent-insoluble fraction and that blocking *de novo* RP production, either by anchoring away Ifh1 or treating cells with cycloheximide, diminishes or abolishes a key transcriptional consequence of RAStR, namely up-regulation of Hsf1 target genes. Our observations thus strongly suggest that RP aggregates are an important activating signal for RAStR. Nevertheless, we and others (Sung et al., 2016a) detect a large number of additional proteins that accumulate in an insoluble fraction upon ribosome assembly stress, including many RiBi proteins (e.g. numerous rRNA helicases and processing factors), the RP gene activator Ifh1 and chaperones. At present we do not know the precise molecular nature of these aggregates, which presumably accumulate in the nucleolar space, or even whether they represent a common structure. In any event, our data suggest that RP- and Ifh1-containing aggregates are highly dynamic since the sequestration of Ifh1 and activation of Hsf1 by Top1 degradation are rapidly reversed by Top2 compensation. We imagine that this provides a strong selective advantage by allowing cells to rapidly recover from a transient disruption of ribosome biogenesis.

Accumulation of proteins in insoluble fractions or aggregates underlies numerous diseases, as well as aging (Saarikangas and Barral, 2015; Tuite and Melki, 2007). However, certain protein aggregates appear to be dynamic structures that contribute to cellular fitness by protecting the cell during stress (Cherkasov et al., 2013; Douglas et al., 2008; Grousl et al., 2018; Kaganovich et al., 2008; Miller et al., 2015b). For example, stress granules and P-bodies, two of the most intensively studied insoluble macromolecular aggregates, have emerged as important cytoplasmic regulators of gene expression by controlling the processing, sequestering and/or degradation of specific RNA transcripts (Decker and Parker, 2012; Mahboubi and Stochaj, 2017). Interestingly, it has recently been reported that nucleolar proteins can form sub-compartmental structures by promoting liquid-liquid phase separation (Berry et al., 2015; Feric et al., 2016). Although further work will be required to characterize the composition, assembly and function of these nucleolar membrane-less structures, they are attractive candidates for regulatory hubs that could act by sensing ribosome biogenesis stress and controlling adaptive responses. With respect to the present study, we imagine that liquid phase-separated structures in the nucleolus could be directly involved in sequestration of Ifh1 during RAStR, as well as the titration of Hsp70 that we propose leads to Hsf1 activation. A challenge for future studies will be to characterize the physical properties of these postulated structures and their relevance to the transcriptional outputs that we measure here.

We showed in a previous study (Albert et al., 2016) that in strains where RNAPI is constitutively active (Laferte et al., 2006) Ifh1 is released from RP gene promoters shortly after TORC1 inhibition by rapamycin treatment (~5 minutes) but returns only 15 minutes later. This promoter re-binding does not occur in wild type cells due to the action of two RiBi proteins (Utp22 and Rrp7) that can sequester Ifh1 in the CURI (CK2/Utp22/Rrp7/Ifh1) complex (Albert et al., 2016; Rudra et al., 2007). The findings reported here suggest that *S. cerevisiae* has developed a second, temporally-separated mechanism that helps to align RP gene expression with ribosome assembly. We propose that this short timescale mechanism results from a rapid rise in unassembled RPs that occurs immediately following TORC1 inhibition, or other stresses that disrupt ribosome assembly, such as depletion of Top1 or RiBi factors. Notably, we show that the Hsf1 regulon is also differentially regulated at short and long timescales following TORC1 inactivation. We propose that the rapid induction of Hsf1 target genes in combination with arrest of RP gene expression caused by RAStR induction contributes to the clearing of proteotoxic unassembled RPs from the nucleus, thereby removing the signal for Hsf1 activation. The long timescale mechanism described previously allows the cells to resume RP transcription in coordination with RNAPI activity, in order to reinstate a balance in the production of ribosomal precursors.

As alluded to above, ribosome assembly stress in higher eukaryotes has been studied extensively in the context of ribosomopathies, diseases often associated with RP gene haplo-insufficiencies, RP gene point mutations or mutations in RiBi factors. One hypothesis put forward to explain these observations is that unassembled RPs trigger a feedback mechanism that decreases transcription of ribosome biogenesis genes by inhibiting c-Myc function and arrests cell growth through p53 activation (Dai et al., 2007; Liu et al., 2016). It has also been recently reported that the rRNA helicase DDX21 binds to and activates RP gene promoters in a manner that may be sensitive to the status of ribosome biogenesis (Calo et al., 2015). Taken together, these findings suggest that unassembled RPs could mediate an ancestral process to regulate ribosome biogenesis conserved from prokaryotes (Nomura, 1999) to eukaryotes. Transcriptome analysis immediately following ribosome assembly stress in mammalian cells will be required to understand the interplay between these different mechanisms and may also uncover novel pathways.

Our work also provides insights into the connection between ribosome assembly and Hsf1 first revealed in a report from the Churchman lab that appeared as our work was being prepared for publication (http://dx.doi.org/10.1101/458810). Hsf1 is a key sensor of proteotoxic stress in all eukaryotes that controls a common set of chaperones conserved from yeast to human. One protective function reported for Hsf1 is its ability to reduce protein aggregate formation leading to neurogenerative diseases (Neef et al., 2014). On the other hand, Hsf1 also exerts a pro-oncogenic function through its ability to promote proteostasis in rapidly growing tumor cells (Mendillo et al., 2012; Santagata et al., 2011). Despite its central function, a holistic understanding of the regulatory mechanisms that govern Hsf1 activity still missing. Our work and that of Tye et al. (http://dx.doi.org/10.1101/458810) demonstrates that Hsf1 activity is tightly linked to ribosome biogenesis in yeast, in a manner independent of the previously described RQC mechanism that contributes to the dissociation of aberrant nascent polypeptides from the ribosome (Brandman et al., 2012). These two mechanisms highlight the central importance of ribosome assembly and activity in regulation of cellular protein homeostasis through Hsf1. Although it is currently unknown if RAStR is conserved in metazoans, we note that RPs are also subjected to a high turnover rate compared to other nuclear components in mammalian cells and that proteasome or ribosome assembly inhibition trigger a rapid accumulation of RPs in the nucleus, whereas arrest of translation has an opposite effect (Lam et al., 2007; Sung et al., 2016a). Importantly, it was reported that cycloheximide treatment also abolishes Hsf1 activity in mammalian cell by an unknown mechanism (Santagata et al., 2013). We propose that a dynamic balance between unassembled and assembled RPs could be sensed by Hsf1 to constantly adjust protein homeostasis transcription programs in eukaryotes with translational flux, proteolysis and the rate of ribosome assembly (**Figure 7**), since disruption or hyperactivation of any of these processes will rapidly change nuclear levels of free RPs. Given the growing body of evidence linking Hsf1 activity to numerous diseases associated with proteotoxic stress, but also rapid cell growth in cancer, it will be of great interest to challenge this model in the future.

## Materials and Methods

### ChIP-Seq

Cultures of 50 mL in YPAD were collected at OD_600_ 0.4 - 0.8 for each condition. The cells were crosslinked with 1% formaldehyde for 5 min at room temperature and quenched by adding 125 mM glycine for 5 min at room temp. Cells were washed with ice-cold HBS and resuspended in 3.6 mL of ChIP lysis buffer (50 mM HEPES-Na pH 7.5, 140 mM NaCl, 1mM EDTA, 1% NP-40, 0.1% sodium deoxycholate) supplemented with 1mM PMSF and 1x protease inhibitor cocktail (Roche). Samples were aliquoted in Eppendorf tubes and frozen. After thawing, the cells were broken using Zirconia/Silica beads (BioSpec). The lysate was spun at 13,000 rpm for 30 min at 4°C and the pellet was resuspended in 300 μl ChIP lysis buffer + 1mM PMSF and sonicated for 15 min (30 sec ON - 60 sec OFF) in a Bioruptor (Diagenode). The lysate was spun at 7000 rpm for 15 min at 4°C. Antibody (1 μg / 300 μL of lysate, Abcam ab5131) was added to the supernatant and incubated for 1 hr at 4°C. Magnetic beads were washed three times with PBS plus 0.5% BSA and added to the lysates (30 μL of beads/300 μL of lysate). The samples were incubated for 2 hr at 4°C. The beads were washed twice with (50 mM HEPES-Na pH 7.5, 140 mM NaCl, 1mM EDTA, 0.03% SDS), once with AT2 buffer (50 mM HEPES-Na pH 7.5, 1 M NaCl, 1mM EDTA), once with AT3 buffer (20 mM Tris-Cl pH 7.5, 250 mM LiCl, 1mM EDTA, 0.5% NP-40, 0.5% sodium deoxycholate) and twice with TE. The chromatin was eluted from the beads by resuspension in TE + 1% SDS and incubation at 65°C for 10 min. The eluate was transferred to an Eppendorf tube and incubated overnight at 65°C to reverse the crosslinks. The DNA was purified using High Pure PCR Cleanup Micro Kit (Roche). DNA libraries were prepared using TruSeq ChIP Sample Preparation Kit (Illumina) according to manufacturer’s instructions. The libraries were sequenced using an Illumina HiSeq 2500 and the reads were mapped to sacCer3 genome assembly using HTSStation (shift = 150 bp, extension = 50 bp; (David et al., 2014)). For ChIP-qPCR, primer sequences are available upon request. To compare depleted versus non-depleted cells, we divided the signal from the +auxin and/or rapamycin and/or cycloheximide sample by the signal from the – auxin and/or rapamycin and/or cycloheximide (vehicle) sample and log2 transformed this value. All data from publicly available databases were mapped using HTS Station (http://htsstation.epfl.ch; (David et al., 2014)).

### Yeast strains and growth

Strains used in this study are listed in **Table S7.**Experiments were typically performed with log phase cells harvested between OD_600_ 0.4 and 0.8. Anchor-away of FRB-tagged proteins was induced by the addition of rapamycin (1 mg/ml of 90% ethanol/10% Tween stock solution) to a final concentration of 1 μg/ml (Haruki et al., 2008). Depletion of AID-tagged protein was induced by the addition of auxin (3-indoloacetic acid) at 500 μM final concentration. Arrest of translation was induced by the addition of cycloheximide to a final concentration of 25 μg/ml. Cells are treated with diazaborine to a final concentration of 50ug/ml.

### Fluorescence microscopy

Cells were grown overnight at 30°C in SC medium (0.67% nitrogen base without amino acids (BD), 2% dextrose supplemented with amino acids mixture (AA mixture; Bio101), adenine, and uracil). Cells were diluted and were harvested when OD_600_ reached 0.4. Cells were spread on slides coated with an SC medium patch containing 2% agarose and 2% glucose. Stacked images were recorded (Intelligent Imaging Innovations) at a spinning disc confocal inverted microscope (Leica DMIRE2; Leica) using the 100x oil objective and an Evolve EMCCD Camera (Photometrics).

### Insoluble fraction purification and mass spectrometry

Isolation of protein aggregates from yeast cells was performed as described previously (Koplin et al., 2010) with slight modifications. 50 OD_600_ units (50 ml) of exponentially growing cells were harvested, and cell pellets were frozen in liquid N2. The cell pellets were resuspended in 1 ml lysis buffer (20 mM Na-phosphate pH 6.8, 10 mM DTT, 1 mM EDTA, 0.1% Tween, 1 mM PMSF, protease inhibitor cocktail and 100 units/ml zymolyase) and incubated at 30° C for 30 min. Chilled samples were treated by tip sonication (20%, 10 sec, 2x) and centrifuged for 20 min at 600 g at 4°C. Aggregated proteins were pelleted at 16,000 g for 20 min at 4°C. After removing supernatants, insoluble proteins were washed once with Wash I buffer (20 mM Na-phosphate pH 6.8, 500 mM NaCl, 5 mM EDTA, 2% NP-40, 1 mM PMSF, and protease inhibitor cocktail), and centrifuged at 16,000 g for 20 min at 4°C. Insoluble proteins were washed with Wash II buffer (20 mM Na-phosphate pH 6.8(Cold)), pelleted and sonicated (10%, 10 s, twice) in 40 μl of Wash II buffer. For analysis by SDS-PAGE (4–12% acrylamide) and subsequent Western blotting, proteins were first boiled in Laemmli buffer. 1x of the total cell lysate (T) and 20x of the isolated pellet fraction (P) were separated and analyzed by Coomassie Blue staining or immunoblotting. Proteins were identified by shotgun mass spectrometry analysis at the Functional Genomics Center Zurich (ETH, Zurich) following TCA precipitation (20%) and acetone washing, according to posted procedures. Database searches were performed by using the Mascot (SwissProt, all species; SwissProt, yeast) search program, using very stringent settings in Scaffold (1% protein FDR, a minimum of 2 peptides per protein, 0.1% peptide FDR).

### Polysome gradients

Yeast cells growing exponentially were treated or not with auxin for 20 min. 50 μg/ml cycloheximide (Sigma) is added directly to the culture medium. Cells were collected by centrifugation, rinsed with buffer K [20 mM Tris-HCl pH 7.4, 50 mM KCl, 10 mM MgCl_2_] supplemented with 50 μg/ml cycloheximide and collected again by centrifugation. Dry pellets were resuspended with approximately one volume of ice-cold buffer K supplemented with 1 mM DTT, 1× Complete EDTA-free protease inhibitor cocktail (Roche), 0.1 U/μl RNasin (Promega) and 50 μg/ml cycloheximide. About 250 μl of ice-cold glass beads (Sigma) were added to 500 μl aliquots of the resuspended cells and cells were broken by vigorous shaking, 3 times 2 min, separated by 2 min incubations on ice. Extracts were clarified through 2 successive centrifugations at 13 000 rpm and 4°C for 5 min and quantified by measuring absorbance at 260 nm. About 30 A260 units were loaded onto 10% - 50% sucrose gradients in buffer K, and then centrifuged for 150 min at 39,000 rpm and 4°C in an Optima L-100XP Ultracentrifuge (Beckman-Coulter) using a SW41Ti rotor without brake. Following centrifugation, 18 fractions of 500 μl each were collected from the top of the gradients with the Foxy Jr. apparatus (Teledyne ISCO). The absorbance at 254 nm was measured during collection with the UA-6 device (Teledyne ISCO).

### Pulse labeling, RNA extraction and Northern hybridization

Metabolic labeling of pre-rRNAs was performed as previously described (Tollervey et al., 1993) with the following modifications. Strains were grown in synthetic glucose medium lacking adenine to an OD_600_ of 0.8. Auxin (0.5mM) was next added to the cultures and cells were labeled for 2 min with [2,8-^3^H]-adenine (NET06300 Perkin Elmer) at 0, 10, 20 and 30 min following the addition of auxin. Cell pellets were frozen in liquid nitrogen. RNA extractions and Northern hybridizations were performed as previously described (Beltrame and Tollervey, 1992). For high molecular weight RNA analysis, 2 μg of total RNA were glyoxal denatured and resolved on a 1.2% agarose gel. Note that Northern hybridization was performed on [2,8-^3^H]-adenine labeled RNA. Membrane was first exposed to reveal neo-synthetized transcripts, and subsequent Northern hybridization revealed rRNA transcript abundance.

## Acknowledgments

We would like to thank Mylène Docquier and the Genomics Platform of iGE3 at the University of Geneva (https://ige3.genomics.unige.ch/) for high throughput sequencing services, the Functional Genomics Center Zurich (ETH, Zurich) for mass spectrometry analysis, Prof. Helmut Bergler (Karl-Franzens-Universität, Graz, Austria) for his generous gift of diazaborine, Lyudmil Raykov and the Bioimaging Center at the Faculty of Sciences, University of Geneva (http://bioimaging.unige.ch/) for help with confocal microscopy, Nicolas Roggli for expert assistance with data presentation and artwork, and all members of the Shore lab for comments and discussions throughout the course of this work. B.A. acknowledges support from a long-term EMBO postdoctoral fellowship in the early phases of this work. M.J.B. was supported in part by an iGE3 Ph.D. student fellowship. D.S. acknowledges funding from the Swiss National Science Foundation (grant number 31003A_170153) and the Republic and Canton of Geneva.

## Data availability

Read counts for all RNAPII ChIP-seq experiments (integrated counts over the complete open reading frame of all protein-coding genes) are given in Supplementary Tables (**Tables S1-6**). Primary processed sequence files will be made available at Gene Expression Omnibus (GEO accession number pending).

