## Supplemental Figures S1-S5 for "A ribosome assembly stress response regulates transcription to maintain proteome homeostasis"

### Supplemental Figure S1

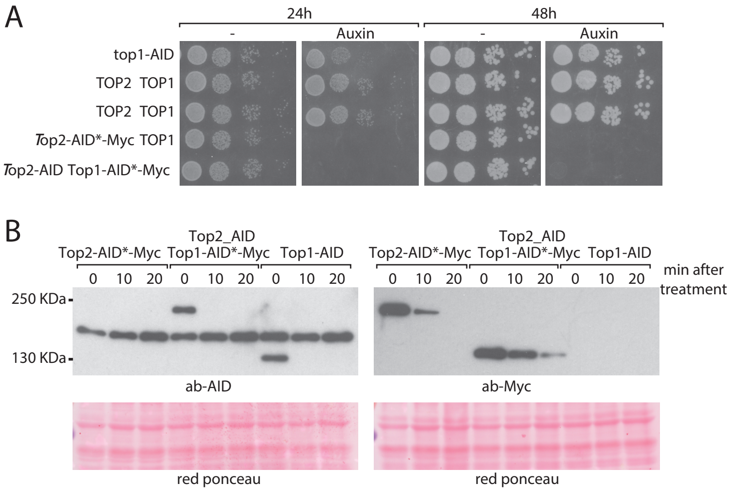


C

| GO term | p-values |
|---|---|
| protein folding | 1.71E-10 |
| protein refolding | 3.39E-07 |
| response to heat | 0.00141178 |
| response to temperature stimulus | 0.00235991 |
| response to abiotic stimulus | 0.02935877 |

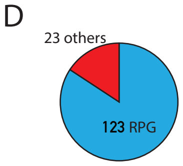


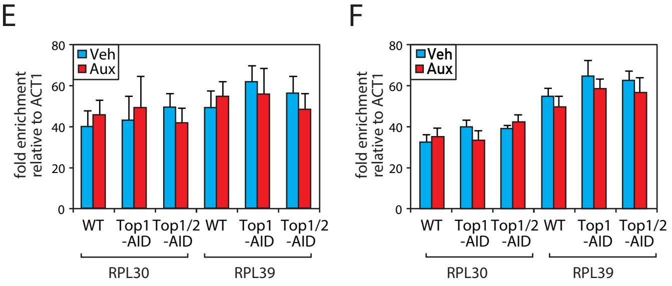


**Supplemental Figure S1** (related to Figure 1): (**A**) 10-fold serial dilution of Top1-AID or Top1/2-AID cells spotted onto rich medium and grown at 30°C for 24 or 48 hours with or without auxin treatment. (**B**) Top1 or Top2 tagged as indicated were detected by western blot using antibodies against Myc or AID antibody in a whole cell extract. Ponceau red stain served as a loading control (bottom panel). (**C**) GO of genes that are upregulated (Rpb1-ChIP) more than two times following Top1 depletion. (**D**) Circle diagram showing number of RP genes in the 146 genes downregulated more than two times following Top1 depletion. (**E-F**) Rap1 (E) or Fhl1 (F) occupancy at the RPL30 and RPL39 promoters at 20 min following Top1 or Top1/2 depletion.

### Supplemental Figure S2

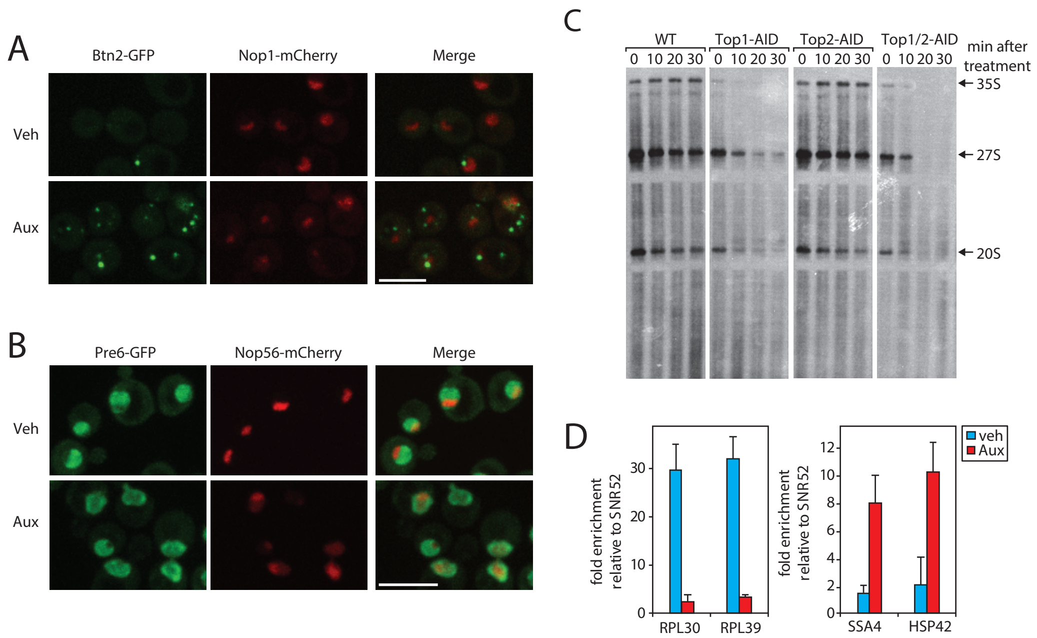


**Supplemental Figure S2** (related to Figure 2): (**A-B**) Yeast strain expressing Btn2-eGFP (A) or Pre6-eGFP (B) and Nop1-mCherry was grown exponentially and cell samples were used for fluorescence microscopy analysis after 20 min of Top1/2 depletion by auxin treatment (Aux) or without depletion (Veh). Scale bars correspond to 2 µm. (**C**) Yeast cells were grown to mid-log phase and auxin was added within the media and cells were then pulse labeled with [3H] adenine for 2 min aliquots at different time points after Auxin addition (T=0, 10, 20, 30 min) in WT, Top1-AID, Top2-AID, and Top1/2-AID strains. RNAs were extracted and samples were separated on agarose gels, transferred to a nylon membrane. (**D**) RNAPII ChIP occupancy at the *RPL30* and *RPL39* promoters or at the *SSA4* and *HSP42* promoters at 20 minutes following Top1 and Top2 depletion.

### Supplemental Figure S3

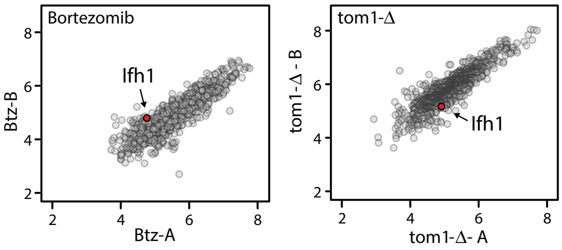


**Supplemental Figure S3** (related to Figure 3): Scatter plots representing ΔiBAQ of biological replicate B (y-axis) versus replicate A (x-axis) for insoluble proteins in btz-treated cells (left panel) or in *tom1*-Δ cells (right panel). Red spots indicate Ifh1. Raw data are from (Sung et al., 2016a; Sung et al., 2016b).

### Supplemental Figure S4

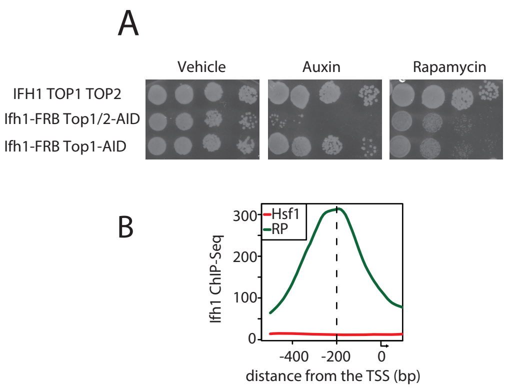


**Supplemental Figure S4** (related to Figure 4): (**A**) 10-fold serial dilution of cells with the indicated *IFH1*, *TOP1* and *TOP2* alleles were spotted onto rich medium containing either vehicle (control), auxin, or rapamycin, as indicated, and grown at 30°C for 40 hours before being photographed. All strains are of the rapamycin-resistant (nuclear) anchor-away background (*TOR1-1 RPL13A-FKB12*). (**B**) Average Ifh1 ChIP-seq signal with respect to the TSS at promoters of RP genes (green) or Hsf1 target genes (red).

Supplemental Figure S5

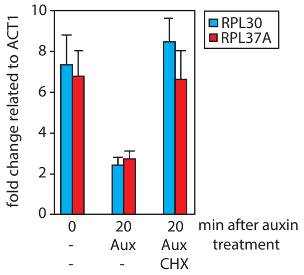


**Supplemental Figure S5** (related to Figure 5): Ifh1 occupancy at the RPL30 and RPL37A promoters at 0 or 20 min following Top1-AID and Top1/2-AID depletion (Aux) in presence (CHX) or absence (-) of cyclohex-imide.
